# The Promotive Effect of Ocean Literacy on Marine Conservation Behavior: A Qualitative Study Based on Chinese University Students

**DOI:** 10.1101/2025.04.11.648364

**Authors:** Linzhao Wang, Bo Gao, Xiangwei Chang, Li Zhang

## Abstract

**Introduction:** Ocean literacy is crucial in shaping marine conservation behaviors among Chinese university students. This study explores the intricate relationship between ocean literacy and pro-environmental behaviors, emphasizing the roles of emotional connections, cultural significance, and educational experiences. By examining the multidimensional aspects of ocean literacy, the research seeks to identify key factors that influence conservation actions and the practical application of marine knowledge.

**Methods:** Utilizing qualitative methodologies, this study conducted semi-structured interviews with 16 Chinese university students to delve into their perceptions and interactions with the marine environment. The interviews explored various dimensions such as oceanic knowledge, environmental beliefs, value orientations, and actions related to marine conservation. NVivo software was used for data analysis, allowing for the identification of key themes through a rigorous coding process. The study’s analysis also included a moderated mediation model to examine how educational experiences influence the relationship between ocean literacy and conservation behaviors.

**Results:** The findings reveal that ocean literacy among university students comprises key components such as scientific knowledge, environmental ethics, conservation action, policy understanding, and critical thinking. Significant gaps in marine knowledge were observed, alongside strong emotional and cultural connections to the ocean, which significantly mediate the relationship between ocean literacy and conservation behaviors. Educational experiences further moderated these effects, highlighting the need for enhanced marine education.

**Conclusion:** This study underscores the necessity of integrating comprehensive educational strategies with emotional and cultural engagement to effectively promote marine conservation behaviors. The results advocate for educational reforms that incorporate practical and interactive learning approaches, aimed at deepening students’ understanding and involvement in marine conservation. Strengthening these educational frameworks can significantly enhance ocean literacy and foster proactive environmental stewardship among university students.

## Introduction

The ocean, as Earth’s largest life-support system, plays a critical role not only as a reservoir of biodiversity but also as a regulator of global climate systems. Marine ecosystems, including coral reefs, seagrass beds, and mangrove forests, provide essential ecosystem services such as carbon sequestration, oxygen production, and habitats for millions of species [1]. The ocean’s health is indispensable for global ecological equilibrium and the sustainable development of human civilization. However, anthropogenic activities, such as overfishing, pollution, and habitat destruction, increasingly threaten marine ecosystems, adversely impacting biodiversity and jeopardizing livelihoods globally [2].

In response to the escalating oceanic ecological crises, marine protection has become a focal point on the global environmental agenda. The United Nations (UN) and World Health Organization (WHO) have called for interdisciplinary and multi-sectoral collaboration at all governance levels to strengthen human-nature connections and enhance both human well-being and environmentally responsible behaviors [3]. Oceans are central to the UN Sustainable Development Goals (SDGs), linking the environmental, social, and economic pillars of sustainability. Initiatives such as the “Decade of Ocean Science for Sustainable Development” emphasize the importance of capacity building, technology transfer, education, and international cooperation in marine science to improve global marine health [4]. Furthermore, international treaties, including the United Nations Convention on the Law of the Sea (UNCLOS) and the Convention on Biological Diversity (CBD), underscore international responsibilities and cooperation in ocean conservation efforts.

Marine education, notably ocean literacy, plays a pivotal role in fostering environmental protection and sustainable development. Originating in the United States in 2002, the concept of Ocean Literacy (OL) was introduced to improve public understanding of ocean issues and promote responsible actions toward ocean sustainability [5]. Fauville articulated ocean literacy as encompassing knowledge of fundamental marine concepts, effective communication about marine issues, and informed decision-making and responsible actions regarding ocean resources [6]. Recent scholarship has expanded the concept of ocean literacy beyond basic marine science knowledge, incorporating attitudes, behaviors, and the emotional connections individuals have with the ocean, which are critical for sustainable ocean management [7, 8].

Enhancing ocean literacy has emerged as a key strategy for addressing marine sustainability challenges, particularly by engaging youth—who represent a crucial demographic in global efforts to mitigate climate change and ocean degradation [9]. Indeed, evidence suggests that increased ocean literacy positively influences environmental behaviors, leading to more proactive and responsible participation in ocean conservation efforts [10]. However, this relationship is complex and shaped by psychological mediators such as environmental beliefs, values, and a sense of responsibility. Frameworks such as the Theory of Planned Behavior (TPB), the Value-Belief-Norm (VBN) theory, and the Norm Activation Model (NAM) have been instrumental in explicating the psychological pathways that transform environmental awareness into tangible pro-environmental actions [11-13].

Despite growing global attention to ocean literacy, research within the Asian context, particularly among Chinese university students, remains limited. While studies in Western contexts are relatively abundant, understanding in the Asian context—especially within China—is notably lacking [14, 15]. Given China’s extensive coastline spanning over 14,000 kilometers and the pressing threats posed by marine pollution, overfishing, and ecological degradation, examining ocean literacy among Chinese university students is critical. Enhancing ocean literacy among this demographic not only addresses immediate environmental challenges but also builds long-term capacity for sustainable marine management.

This study aims to bridge existing gaps by qualitatively exploring the mechanisms through which ocean literacy fosters pro-environmental behaviors among Chinese university students. Specifically, we investigate how ocean literacy interacts with environmental beliefs, values, and the sense of responsibility to influence marine environmental behaviors. By providing empirical insights into these complex psychological relationships, the research seeks to inform targeted educational strategies and policy interventions that effectively promote marine sustainability in China and beyond.

### Theoretical Framework

Ocean literacy serves as a multidimensional construct underpinned by interdisciplinary theories drawn from environmental psychology, education, and sustainability science. Cava et al. initially defined ocean literacy as “the understanding of the ocean’s influence on individuals and society, and vice versa, the influence of individuals and society on the ocean” [5]. They posited that individuals with a high level of ocean literacy should possess three key capabilities: 1) understanding of fundamental oceanic knowledge; 2) the ability to communicate information about the ocean in meaningful ways; and 3) the capacity to make informed and responsible decisions when confronted with issues concerning the ocean and its resources [6]. Additionally, the European Marine Board in 2013 emphasized that ocean literacy primarily refers to people’s cognition and understanding of the ocean, enabling them to make wise decisions between marine environments and human activities, and to take actions at both individual and collective levels to support the sustainable development and future protection of the ocean [16].Fundamentally, ocean literacy involves understanding essential principles of marine ecosystems, human impacts on oceans, and the reciprocal relationships between humans and marine environments [5, 6]. This comprehensive understanding extends beyond knowledge acquisition, encompassing attitudes, emotional connections, and behaviors that foster marine stewardship and sustainable ocean management [7, 8].

The theoretical foundation guiding this qualitative investigation integrates three prominent theoretical models: the Theory of Planned Behavior (TPB) emphasizes the critical role of attitudes, subjective norms, and perceived behavioral control in shaping individual intentions and subsequent behaviors. In the context of ocean literacy, TPB implies that enriched marine knowledge and positive attitudes towards ocean conservation significantly influence individuals’ behavioral intentions and ultimately their conservation behaviors [11]. The Value-Belief-Norm theory (VBN), proposed by Stern, further elaborates on the interplay among personal values, ecological worldviews, and normative beliefs in influencing environmental actions [17]. Specifically, VBN emphasizes that when individuals possess strong biospheric and altruistic values, they develop heightened environmental awareness and perceived responsibility, resulting in greater engagement in pro-environmental behaviors. Within this framework, ocean literacy enhances individuals’ ecological consciousness and fosters a heightened sense of responsibility towards marine ecosystems, prompting responsible and sustained conservation behaviors. Complementing the TPB and VBN theories, the Norm Activation Model (NAM) introduced by Schwartz highlights the critical role of moral obligations and perceived environmental responsibility in activating pro-environmental behaviors [18]. NAM posits that individuals become more inclined to engage in environmentally beneficial actions when they recognize environmental threats and perceive a personal responsibility to mitigate such threats. Ocean literacy, therefore, can serve as a catalyst that activates moral responsibility toward marine conservation by enhancing awareness of the ocean’s critical condition and personal accountability in addressing marine issues.

Moreover, qualitative research methods offer a unique opportunity to deeply explore individual experiences, beliefs, and emotional connections with the marine environment. By employing qualitative methodologies, such as semi-structured interviews and thematic analysis, this study can uncover nuanced insights into how ocean literacy affects the attitudes and behaviors of Chinese university students. The qualitative approach is particularly suited to capture complex, culturally specific interpretations of ocean-related environmental behaviors, which quantitative methods alone may overlook [19, 20].

In summary, this theoretical framework, anchored in TPB, VBN, and NAM, combined with qualitative methodologies, provides a robust basis for understanding the complex mechanisms through which ocean literacy influences marine environmental behaviors among Chinese university students.

### Research Status and Hypothesis Development

Recent developments in ocean literacy research have expanded the concept from an exclusively knowledge-based model to a broader framework that integrates attitudes, behaviors, emotional connections, and individual engagement with ocean-related issues [7, 8]. Brennan emphasized the multidimensional nature of ocean literacy, outlining critical dimensions such as knowledge, awareness, attitudes, communication, behaviors, and activism, all directly influencing practical marine conservation actions[7]. Similarly, McKinley highlighted the importance of understanding human-ocean relationships in shaping sustainable marine behaviors [8].

The connection between ocean literacy and pro-environmental behaviors is well-documented. Ashley et argued that elevated ocean literacy positively correlates with increased proactive marine environmental actions by raising awareness and triggering responsible decision-making [10]. Existing research also confirms a significant correlation between the level of ocean literacy among university students and their environmental concerns. Surveys in countries like Canada have shown that, despite varying levels of marine science knowledge, students with higher ocean literacy typically demonstrate greater interest in marine environmental protection and sustainable management [21, 22]. This correlation is not limited to student populations but is also prevalent among individuals of different ages, regions, and educational backgrounds [23]. Furthermore, studies have shown that enhancing ocean literacy can significantly strengthen these psychological traits, thereby promoting pro-ocean behaviors [24]. Ocean literacy does not always lead directly to behavioral change but is often influenced by personal environmental beliefs, social norms, and perceived behavioral control [25, 26].

Drawing upon this body of literature, this study proposes several qualitative hypotheses:

H1: Students articulate their understanding of the connection between ocean literacy and pro-environmental behaviors through personal experiences and social influences.

H2: Students’ expressions of ocean values are positively correlated with their awareness and understanding of global and regional oceanic environmental issues.

H3: Education serves as a crucial catalyst in reinforcing students’ sense of environmental responsibility and their willingness to engage in ocean conservation behaviors.

H4: Enhanced ocean literacy enables students to clearly articulate the role and importance of ocean-related beliefs, values, and responsibility in their pro-environmental behaviors.

H5: Students’ ability to convert ocean literacy into specific environmental protection actions is influenced by cultural, social, and customary factors.

By exploring these hypotheses, the present study aims to deepen the understanding of the intricate relationships between ocean literacy, environmental beliefs, values, responsibility, and pro-environmental behaviors among Chinese university students, providing a robust basis for formulating targeted educational and policy interventions.

## Methods

This study aims to explore the level of ocean literacy among Chinese university students and its relationship with pro-environmental behaviors using semi-structured interviews as a qualitative research method to gather in-depth qualitative data.

### Research Design

The research adopts a qualitative design, utilizing semi-structured interviews to comprehensively explore university students’ ocean literacy and its influence on pro-ocean behaviors. The interviews cover multiple dimensions such as oceanic knowledge, environmental beliefs, value orientations, sense of responsibility, and specific actions, aiming to understand in depth how university students form and practice marine environmental consciousness and behaviors.

### Research Participants

Participants were selected using purposive sampling, primarily consisting of Chinese university students. These students have varying levels of marine-related educational experiences or practical involvement and have shown some interest or enthusiasm in marine environmental issues. The study adheres strictly to ethical standards; all participants volunteered for the interviews and signed informed consent forms. Additionally, the study received formal approval from an institutional ethics committee (Approval Number: 1041386-202407-HR-106-02) to ensure ethical compliance in the research process. All interview participants signed informed consent forms before the interviews to ensure the ethical integrity of the study.

### Data Collection

Interviews and data collection began on July 15, 2024 and ended on July 25, 2024, with interviews conducted one-on-one, lasting approximately 30 to 60 minutes each. The interviews addressed students’ ocean literacy, the influence of family, school, and community on their literacy development, the formation process of their marine environmental beliefs, the relationship between these beliefs and daily decision-making behaviors, and actual pro-ocean behaviors and their motivations. All interviews were recorded and transcribed verbatim to ensure the authenticity and completeness of the data.

### Data Analysis

Data analysis was conducted using NVivo software to ensure systematic and rigorous processing. The analysis involved familiarizing with the data, coding, theme extraction, and theoretical interpretation. Initially, open coding was performed to generate initial codes; this was followed by axial coding, where related nodes were integrated to form major themes; finally, selective coding was conducted to identify core theoretical nodes that clearly explain the relationship between university students’ ocean literacy and environmental behaviors. Through this process, eight key themes were identified, including human-ocean interactions, marine environmental values, environmental beliefs and behaviors, socio-cultural influences, policy and resource management strategies, among others. The results of the data analysis not only enriched the theoretical framework of quantitative research but also provided a deep qualitative perspective on how Chinese university students’ ocean literacy can effectively translate into specific environmental protection actions.

### Data Analysis Tool

The study employed NVivo software for the qualitative data analysis, which efficiently assists researchers in systematically organizing, analyzing, and identifying key nodes and their relationships within the interview texts.

Through the aforementioned research design and methods, this study aims to deeply analyze the mechanisms between ocean literacy and environmental behaviors among Chinese university students, providing empirical foundations and strategic insights for effective marine conservation education.

## Results

### Description of Participants

This research selected 16 Chinese university students through random sampling to participate in semi-structured interviews, aiming to gain an in-depth understanding of their personal insights and experiences on the topic of ocean literacy. The interviews were conducted face-to-face, each lasting about 30 minutes to ensure participants had ample time to fully express their views and experiences. The demographic characteristics of the respondents included diverse genders, ages, and academic backgrounds to achieve a representative and varied sample, ensuring a comprehensive understanding of the research topic.

During data collection, all interviews were audio-recorded and subsequently transcribed into text using professional software. The transcribed content totaled 116,962 words, with the longest single interview record being 12,426 words and the shortest being 4,685 words. All texts were meticulously proofread after transcription to ensure the accuracy and reliability of the data. Additionally, to protect the privacy of the participants, all interview texts were strictly anonymized and identified only by numbers. For detailed information on the interview participants and data statistics, see Table 1.

**Table 1.** Characteristics of participants.

| **No.** | **Gender** | **Major** | **Age** | **Interview time**<br>**(minutes)** |
| --- | --- | --- | --- | --- |
| A1 | Female | Liberal arts | 20 | 44.08 |
| A2 | Female | Liberal arts | 19 | 25.57 |
| A3 | Female | Liberal arts | 20 | 29.02 |
| A4 | Female | Sciences | 23 | 32.20 |
| A5 | Female | Engineering | 21 | 34.13 |
| A6 | Female | Sciences | 22 | 25.26 |
| A7 | Male | Sciences | 22 | 27.26 |
| A8 | Female | Engineering | 19 | 20.27 |
| A9 | Female | Engineering | 19 | 33.14 |
| A10 | Male | Sciences | 21 | 32.21 |
| A11 | Male | Engineering | 21 | 23.01 |
| A12 | Male | Engineering | 21 | 20.19 |
| A13 | Male | Sciences | 21 | 24.02 |
| A14 | Male | Engineering | 21 | 24.49 |
| A15 | Male | Sciences | 22 | 22.14 |
| A16 | Male | Sciences | 22 | 26.46 |

### Data Coding and Processing

This research utilized NVivo 12 software for qualitative analysis of the interview data. NVivo, developed by QSR International, is a professional qualitative research tool that assists researchers in organizing, analyzing, and refining textual data, extracting key themes, and gaining deep insights [19]. To ensure the timeliness and reliability of the data analysis, this study adopted a concurrent interviewing and coding approach. After each interview, researchers promptly organized their notes and converted the audio recordings to text using transcription software. The interview texts and research notes were then systematically organized and coded to prepare for subsequent interviews. Using grounded theory as the analytic approach, the data underwent open coding, axial coding, and selective coding to clearly identify the key factors and theoretical framework influencing the ocean literacy and pro-environmental behaviors of Chinese university students [27].

Additionally, a word frequency analysis was conducted on the texts from all 16 interview participants to reveal the interrelationships among marine environmental beliefs, values, sense of responsibility, and pro-marine behaviors. Specifically, the study performed a word frequency count on the 116,962 words of interview text, setting to display the top 50 most frequent words, and merged synonyms. The resulting word cloud, as shown in Figure 1, highlights terms such as “ocean,” “feeling,” “environment,” “protection,” and “literacy,” with “marine protection” and “marine environment” being particularly prominent, reflecting the respondents’ high level of concern for related issues.

**Figure 1.**
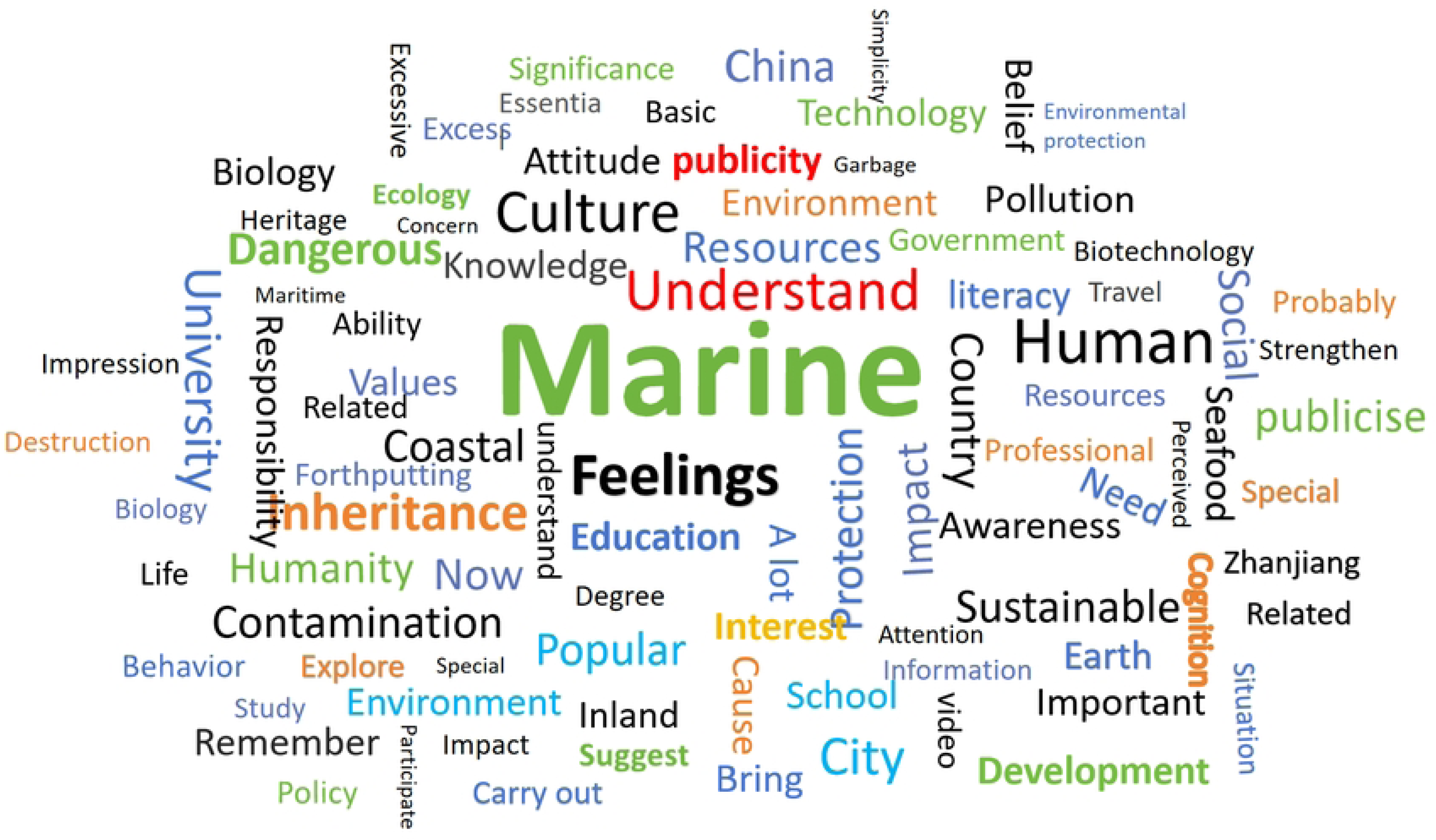

Although the word cloud provides intuitive information, to gain a deeper understanding of the internal structure of university students’ ocean literacy, this study further employed NVivo software for detailed text coding analysis.To thoroughly explore the intrinsic connections between marine environmental beliefs, values, sense of responsibility, and pro-marine behaviors, this study conducted a detailed word frequency analysis of all interview texts. A total of 1001 unique words were identified, with “ocean” being the most frequently occurring word, accounting for 6.05% of the total word frequency. This emphasizes the central position of marine issues in the discussions among university students and aligns with the study’s emphasis on enhancing ocean literacy in environmental education. The frequent appearances of “feeling” (3.41%) and “understanding” (1.15%) reflect the significant roles of emotional connection and cognitive understanding in the interaction process with the ocean in promoting environmental behaviors. Additionally, the presence of key terms such as “protection” (0.97%), “resources” (0.54%), and “development” (0.56%) reveals the students’ concern for the sustainable use and protection of marine resources. During the theoretical linkage analysis, some irrelevant nodes were eliminated, and the top 20 core words by frequency were finally determined, as shown in Table 2.

**Table 2.** Frequency Distribution of Word Occurrence.

| **No.** | **Word** | **Count** | **Weighted**<br>**Percentage (%)** | **Synonyms** |
| --- | --- | --- | --- | --- |
| 1 | Ocean | 1920 | 6.05 | Ocean |
| 2 | Feeling | 1089 | 3.41 | Sensation; Emotion; Contact; Feel;<br>Emotion; Taste; Mood |
| 3 | Understand | 385 | 1.15 | Understand; Comprehend; Recognize; Realize |
| 4 | Protect | 308 | .97 | Preserve; Protect; Guarantee |
| 5 | Culture | 207 | .65 | Culture |
| 6 | Problem | 215 | .65 | Reason Event; Matter; Topic; Problem; |
|  |  |  |  | Subject; Cause; Self |
| 7 | Environment | 210 | .62 | Environment; Situation |
| 8 | Influence | 200 | .60 | Achieve; Arrive; Scope; Region; Effect; Extend; Impact; Function |
| 9 | Important | 179 | .56 | Important |
| 10 | Develop | 180 | .56 | Grow; Develop; Evolve; Develop; Grow |
| 11 | Resources | 171 | .54 | Resources |
| 12 | Utilize | 199 | .52 | Work; Understand; Utilize; Use; Habit; Apply; Master |
| 13 | Place | 174 | .48 | Place; Location; Place; Region; Status; Range; Job; Field; Goal; Area; Position |
| 14 | Country | 138 | .42 | Venue; Country; Territory; Foundation; Land |
| 15 | Literacy | 131 | .41 | Literacy |
| 16 | Feel | 138 | .39 | Hobby; Feel; Experience; Experience; Realization; Experience; Taste |
| 17 | Values | 123 | .39 | Values |
| 18 | Knowledge | 116 | .37 | Scholarship; Knowledge |
| 19 | Humanity | 114 | .36 | Typical; Humanity; Human nature; Someone |
| 20 | Like | 112 | .34 | Hobby; Hope; Like; Willing |

### Final Analysis and Theoretical Node Development

Based on the semi-structured interview outline and relevant ocean literacy theories, we conducted an in-depth comparison, analysis, and integration, ultimately identifying eight main theoretical nodes. These nodes encompass multiple dimensions such as university students’ cognition, emotions, sense of responsibility, and behaviors towards the marine environment, reflecting comprehensively the key factors that influence the formation of ocean literacy among university students. The specific sub-node encodings for each main node are detailed in Table 3. These nodes cover essential areas such as ocean literacy, human-ocean relationships, marine environmental values and beliefs, current environmental challenges, as well as marine responsibility and environmental conservation behaviors. Additionally, the study delved into the socio-cultural impacts of the ocean, marine conservation policies and resource management strategies, and aspects of marine education and public awareness enhancement. These theoretical nodes form a structured analytical framework that reveals the mechanisms of interaction between various dimensions and clarifies how these factors collectively influence both individual and collective ocean literacy cognition and practices.

**Table 3.** Sub-node coding situation table.

| Sub-nodes | Reference Point /<br>Material Text | Subcategory of Reference Point |
| --- | --- | --- |
| Ocean Literacy | 216/16 | Public knowledge of the ocean (32 nodes) |
|  |  | Cognition about the ocean (64 nodes) |
|  |  | Individual ocean literacy levels (52 nodes) |
|  |  | The role and significance of ocean literacy (24 nodes) |
|  |  | Components of ocean literacy (44 nodes) |
| Human-Ocean Relations | 226/16 | The impact of the ocean on humans (50 nodes) |
|  |  | Emotions and attitudes towards the ocean (108 nodes) |
|  |  | Interest and experiences related to the ocean (68 nodes) |
| Marine Environmental Values and Beliefs | 53/16 | Composition of marine values and beliefs (24 nodes) |
|  |  | Impact of marine values and beliefs (29 nodes) |
| Marine Environmental Issues | 105/16 | Societal response to marine pollution (47 nodes) |
|  |  | Public impact of nuclear wastewater incidents (19 nodes) |
|  |  | Impact of marine environmental issues (39 nodes) |
| Marine Responsibility and Behavior | 180/16 | Importance of marine responsibility and conservation behaviors (49 nodes) |
|  |  | Public attitudes and perceptions towards marine conservation (43 nodes) |
|  |  | Participation in marine activities (42 nodes) |
|  |  | Role and significance of marine behaviors (23 nodes) |
|  |  | Challenges and strategies in marine conservation actions (23 nodes) |
| Marine Society and Culture | 73/16 | Importance of marine society and culture (28 nodes) |
|  |  | Protection and inheritance of marine cultural heritage (45 nodes) |
| Marine Policy and Resource Management | 150/16 | Marine policy and international cooperation (43 nodes) |
|  |  | Public perceptions of marine resource utilization (36 nodes) |
|  |  | Marine resource development and conservation (71 nodes) |
| Marine Education and Awareness | 148/16 | Current state of marine education (27 nodes) |
|  |  | Approaches to marine popular science education (43 nodes) |
|  |  | Public awareness of the ocean (49 nodes) |
|  |  | The role and significance of marine education (29 nodes) |

### Interview Research Findings

#### (I) Multidimensional Perspectives on Ocean Literacy

The core components of ocean literacy: Through content analysis of 44 nodes, interview results show that ocean literacy consists of five key elements: scientific knowledge, environmental ethics, conservation action, policy understanding, and critical thinking. These elements work together to influence an individual’s level of ocean literacy. Therefore, strengthening education and practice in these areas is crucial for enhancing the public’s ocean literacy. Public level of ocean knowledge: The public generally lacks systematic ocean knowledge. Although school education covers some marine content, it is often limited to basic concepts of marine biodiversity and ecosystems. This finding is consistent with studies in the Asian region, indicating a lower level of ocean literacy awareness in Asia [21, 28]. Therefore, the existing marine education is insufficient to meet the public’s need for scientific literacy, and promoting ocean literacy education in the Asian region has become an urgent task. Ocean cognition and emotional connection: Respondents have a broad understanding of the physical and biological characteristics of the ocean and its impact on global climate and ecosystems. However, their understanding of deeper environmental issues remains limited. Individual differences in ocean literacy: Analysis has found significant differences in individual levels of ocean literacy. Individuals with high literacy show stronger ocean knowledge and environmental behavior, such as active participation in conservation actions, while those with lower literacy primarily have a basic understanding. Environmental impact of ocean literacy: Ocean literacy is crucial for promoting a sense of environmental responsibility and practical actions. Enhancing public ocean literacy helps increase awareness of the impacts of human activities, driving more effective resource management and the implementation of protective policies [29]. Strengthening marine education plays a central role in enhancing public ocean literacy. Through systematic education and broad social participation, individual and societal awareness of marine environmental issues can be improved, promoting sustainable development.

*A2: “I think it’s quite important, and as a prerequisite for ocean literacy, it’s essential to have a correct understanding. That is, recognizing that the ocean itself is fragile; it is not as strong as it appears.”*

*A11: “The ocean knowledge I’ve encountered is quite limited, and I feel there is also a lack in the spread of this knowledge online. The efforts to popularize it are not sufficient.”*

*A7: “I believe the ocean is the origin of life, and everything on our planet is nurtured from the ocean. So, I see the ocean as the cradle of life for all of humanity and also as a mysterious presence.”*

#### (II) Human-Ocean Relationships

The comprehensive impact of the ocean on humanity: Analysis of 50 coded nodes indicates that respondents generally view the ocean as the origin of life and the frontier of exploration, symbolizing the unknown and freedom. As part of the ecosystem, the ocean has a profound influence on individual self-awareness and sense of environmental responsibility. Ocean-related emotions and attitudes: Based on an analysis of 108 nodes, respondents’ emotions towards the ocean include awe, love, fear, and concern. Many perceive the ocean as a place for relaxation and spiritual healing, and its tranquility is considered beneficial for mental health [7]. Ocean interests and experiences: Respondents commonly express a strong interest in ocean activities (such as swimming, diving, and sailing), believing that direct contact with the ocean deepens emotional connections and enhances awareness of ocean conservation. Interview results highlight the ocean’s importance in promoting mental health, enhancing environmental awareness, and shaping human values [30]. The aesthetic and ecological value of the marine environment also contributes to increasing public pro-environmental behavior [31, 32]. From an evolutionary psychology perspective, human emotional and cognitive connections to the ocean may stem from our evolutionary dependence on water sources [14]. Therefore, ocean education and policy should be leveraged to strengthen public emotional connections to the ocean to promote environmental conservation actions.

*A3: “The ocean is closely linked with our planet. If the ocean is polluted, our Earth’s water resources will also be polluted, which would affect the ecological cycle. This could then have a significant impact on our planet.”*

*A1: “Being enveloped in nature, I feel unified with the water and the wind, and it’s very relaxing. My feelings towards the ocean are definitely fond; I particularly like the ocean.”*

#### (III) Ocean Policy and Resource Management

Ocean policy and international cooperation: Based on 43 coded nodes, the majority of respondents believe that strengthening international cooperation is key to achieving sustainable management of marine resources. Transnational policies, such as the establishment of marine protected areas and the joint management of cross-border resources, are considered important strategies for addressing global challenges such as marine pollution and overfishing [1]. The United Nations’ Sustainable Development Goal 14 specifically emphasizes education for sustainable development of the oceans [33]. Although some countries have implemented relevant policies, respondents still perceive a lag in China’s marine regulations and enforcement, indicating a need for stronger policy implementation. Public attitudes towards the utilization of marine resources: Public attitudes towards the exploitation of marine resources are diverse. On one hand, they are concerned with sustainable resource development; on the other hand, they worry about the impact of overexploitation on ecosystems [34]. Education and awareness raising are considered key to balancing resource utilization with conservation. Marine resource development and protection: Respondents emphasize the importance of marine resources for economic development, such as oil extraction and fisheries. However, they also recognize the ecological risks of overfishing and resource depletion [35]. The European Marine Cooperation Organization has had successful experiences in ocean literacy and resource management, which could serve as a model for China. These organizations promote sustainable marine management through policy formulation, public education, and international cooperation [36].

*A9: “The government and international organizations should take responsibility. Public awareness is often influenced by international and governmental actions. If national governments promote marine conservation, people will fully understand the importance of protecting the oceans, and they will act accordingly. The effort to promote this is quite effective.”*

*A15: “The utilization of marine resources should be planned sensibly, and excessive development should be avoided. Intense exploitation can damage the ecosystem, leading to irreversible consequences.”*

*A7: “Sustainable development is crucial. Each year, there should be a fishing ban period to allow marine life to reproduce more abundantly during this time. This can help restore populations to their previous states and allow us to continue harvesting resources. We should adhere to the principles of sustainable development and protect our marine resources.”*

#### (IV) Ocean Values and Beliefs

The study finds that respondents universally recognize the ocean as an irreplaceable natural resource and emphasize the necessity of protecting marine biodiversity. Respondents focus not only on the direct benefits of the ocean to humanity but also express concerns about the overall health of ecosystems, reflecting ecocentric values. Those with higher levels of education tend to engage more actively in environmental conservation actions, such as participating in ocean cleanup events, supporting no-fishing zone policies, and reducing the use of disposable plastics. Education plays a crucial role in shaping and strengthening ocean conservation values, not only enhancing individual environmental awareness but also increasing their sense of responsibility and willingness to act environmentally. Additionally, social networks significantly influence the formation of ocean values. Some respondents indicate that their environmental actions are influenced by family, friends, or community, highlighting the role of group norms in shaping individual environmental behaviors. Coastal residents, due to their direct contact with the ocean, are more likely to develop a stronger willingness to protect it; in contrast, inland residents tend to have lower levels of concern for the ocean and weaker environmental consciousness. This phenomenon can be explained by the Value-Belief-Norm (VBN) theory, where an individual’s sense of environmental responsibility and values directly influence their pro-environmental behavior [17]. Research results indicate that the internalization process of marine values is crucial for promoting societal-wide environmental behavior. In practice, strengthening marine education, enhancing public environmental responsibility, and leveraging social group influence are key strategies to advance sustainable marine conservation.

*A11: “For marine conservation, it is the belief in the value of the ocean that can restrain oneself from harming it. You must respect, honor, and revere the ocean, and then protect it. Those who have been in contact with the ocean or understand it possess a firmer set of values and beliefs about protecting it.”*

*A13: “Beliefs and values are certainly important; they affect your worldview and, consequently, the actions you take in everyday life. These beliefs and values, when presented internationally, involve actions related to the protection and development of the oceans, ultimately aiming to conserve them.”*

#### (V) Marine Responsibility and Behavior

Importance of marine responsibility and environmental behavior: Based on 49 coded nodes, research indicates that individuals play a crucial role in marine conservation. Although the public has a certain level of environmental awareness, there is still a gap in actual action. Respondents believe that small actions such as reducing the use of disposable plastics and participating in beach cleanups can have a positive impact, and enhancing personal and collective responsibility is an effective path to promote environmental behavior [37]. Public attitudes and participation in marine conservation: Respondents generally have a positive attitude towards marine conservation, but there are still deficiencies at the knowledge and behavior levels. Environmental attitudes, knowledge, and perceptions directly influence environmental actions, and organizing practical activities like beach cleanups is considered an effective strategy to enhance environmental awareness [11, 12]. Participation in marine activities: Participation in marine activities is influenced by geographical location, educational background, and personal interest. Coastal residents are more likely to engage in marine activities, while inland residents have less exposure. Previous research indicates that geographical location affects individual participation in marine behaviors [2]. Significance and challenges of marine behavior: Research finds that marine behavior involves not only ecological protection but is also closely related to social, economic, and cultural aspects. A healthy marine environment promotes the development of tourism and fisheries, and also contributes to human health and welfare. However, marine conservation still faces challenges such as limited resources, inadequate regulations, and low public participation. This study shows that a healthy marine environment is crucial for sustainable societal development. Enhancing conservation effectiveness through multi-party cooperation is essential, and the key to promoting marine environmental behavior lies in strengthening education, policy reform, and public participation [11].

*A2: “If we don’t have a sense of responsibility to protect the ocean, it will get polluted. Even a small action can have a big impact on the ocean.”*

*A8: “The general public’s concern for the ocean is not very high, and some people are not even aware of it, with a lot of pollution still being dumped into the sea. A sense of responsibility is very important, and more people should be encouraged to feel responsible for the ocean; the effort of a few is always not enough.”*

*A3: “I’ve participated in a beach clean-up organized by my school, which I found very meaningful. Sometimes I like walking on the beach, and a couple of days ago, I also went to a shipyard with an activity and went aboard a ship.”*

*A14: “A sense of responsibility towards the marine environment is crucial for ocean protection but faces challenges and issues in practice. It is advisable to start small, focusing on everyone’s voluntary actions. Promoting environmental protection extensively, perhaps by setting up eco-friendly slogans in entertainment venues, could raise everyone’s environmental awareness.”*

#### (VI) Marine Society and Culture

The importance of marine society and culture: Analysis from 28 nodes reveals that the ocean plays an irreplaceable role in the construction of society and culture. Most respondents emphasized the profound impact of the ocean on culture and society, not only reflected in the daily life of coastal areas but also deeply influencing the cultural educational content of inland areas. The geographical environment has a significant impact on the formation and development of culture. Additionally, the ocean serves as an important platform for economic and cultural exchanges; historically, the “Maritime Silk Road” exemplifies its role in global cultural exchanges, facilitating cultural and commodity exchanges between East and West. Marine culture includes not only material culture, such as fisheries and navigation technologies, but also intangible culture, such as traditional customs and religious beliefs. These cultural elements together constitute the uniqueness of marine societies and are significant for national and regional cultural identity. Protection and inheritance of marine cultural heritage: Analysis of 45 nodes indicates that the protection and inheritance of marine cultural heritage are among the key topics discussed by respondents. Many expressed support for the preservation of marine cultural heritage, such as maritime history, fishing village culture, and marine mythology (e.g., Mazu culture). They believe that these cultural heritages are not only a preservation of history but also an educational resource for future generations. However, in the context of rapid modernization, these traditional knowledges and customs face the risk of being forgotten. Cultural heritage management theory suggests that protecting cultural heritage requires consideration of economic, social, and environmental factors. Strengthening the protection of marine cultural heritage not only helps maintain cultural diversity but also contributes to achieving sustainable development goals [38]. This requires the joint efforts of governments, educational institutions, and the public, along with more systematic policy support.

*A3: “It’s quite important in cultural aspects, with many literary works and paintings inspired by the ocean. So, it’s quite significant in the arts.”*

*A5: “It’s definitely worth protecting! It’s also a part of Chinese traditional culture. Some coastal communities have depended on fishing for generations, and these cultural traditions passed down are a valuable heritage.”*

*A6: “I’ve heard about Mazu, and many people believe in her, especially in Fujian. The Maritime Silk Road is itself a cultural heritage that facilitated the exchange between Eastern and Western civilizations, not only allowing for trade but also for the exchange of technologies.”*

#### (VII) Marine Education and Awareness

Current State of Marine Education: Analysis from 27 nodes reveals significant disparities in the prevalence and depth of marine education across different regions and institutions. While some areas have introduced courses related to marine economics, most respondents note that these courses are heavily theoretical, lacking in practicality and interactivity, which limits students’ interest in and deep understanding of marine science [39]. Moreover, marine education urgently needs improvement in content richness, pedagogical methods, and practical applications. Limitations in marine education may lead to students lacking a comprehensive understanding of marine ecosystems, thereby affecting their willingness to participate in marine conservation. Thus, enhancing the quality and effectiveness of marine education has become a pressing issue. Paths to Marine Science Popularization: Analysis of 43 nodes shows that the public primarily acquires marine knowledge through school education, online platforms, social media, and public science popularization activities. Although the internet and social media offer a wealth of marine science content, issues with information fragmentation and superficiality persist. Some respondents particularly emphasize the importance of firsthand experiences, noting that direct contact with the marine environment aids in a profound understanding of marine ecology and the necessity of its protection. Diversified popularization approaches can enhance public attention to marine issues. For instance, venues like marine museums and aquariums provide interactive learning environments that help raise public ocean literacy [40].

*A2: “In my primary, middle, and high school, they only said not to play by the riverbank, not the seaside, and there wasn’t much spread about this. There was also no talk of marine education.”*

*A16: “As a child, I learned about the ocean through books, like ‘A Hundred Thousand Whys’ and other marine science popular books. Later on, I gradually learned more through the internet and science videos, which also delved deeper.”*

Public Awareness of Marine Issues: Interviews reveal that public awareness of marine conservation is generally low, especially among residents of inland areas who are less concerned about marine issues. Most people’s understanding of marine conservation is limited to basic knowledge, lacking in-depth understanding and motivation for action. This indicates that strengthening social outreach and education for marine protection is essential to enhance public understanding of marine ecosystem services and conservation needs. An individual’s environmental attitudes and knowledge directly impact their conservation behaviors, making it crucial to enhance public awareness of marine issues to foster marine conservation actions. Role and Significance of Marine Education: Analysis from the nodes suggests that marine education is crucial for enhancing the awareness of marine conservation, especially among the younger generation. Education not only enhances individual understanding of the importance of marine conservation but also promotes broader environmental protection policies and practices at the societal level. By embedding ongoing conservation behaviors and attitudes through education, marine science knowledge dissemination and application are promoted. Ocean literacy education aims to go beyond traditional knowledge transfer, advocating for diversified learning methods and emphasizing the importance of emotional responses to environmental issues and broader environmental ethics [41]. Previous research indicates the necessity of adopting a more comprehensive approach to ocean literacy, shifting the educational focus from mere knowledge accumulation to transformative marine environmental learning [42].

*A9: “I don’t think it’s very good, many people mainly see the ocean as just beautiful, right? In reality, there isn’t a well-formed, systematic awareness about the ocean. I think the level of ocean awareness among the Chinese public is not sufficient; it’s quite weak.”*

*A6: “There’s a lot of trash, especially on the beaches where many people go to play. You see trash floating around by the seaside, which shows that their ocean awareness is quite weak.”*

#### (VIII) Marine Environmental Issues

Public Response to Marine Pollution: Based on an analysis of 47 nodes, most interviewees expressed significant concern about marine pollution, noting that it directly impacts their participation in marine activities. For example, respondents mentioned that pollution might decrease interest in participating in marine activities, especially when the ocean is perceived as unsafe or unclean. Furthermore, public reactions to marine pollution also extend to economic and health aspects, showing concerns about declines in tourism revenue and long-term health consequences. Respondents generally believe that marine pollution affects not only personal leisure activities but also has negative impacts on community economics and public health. This indicates that the public is aware of the broad harms of marine pollution and recognizes the need for measures to address it. The issue of Japanese nuclear wastewater has triggered widespread concern and discussion among the Chinese public, serving as a catalyst for heightened environmental awareness. Interviews show that such incidents have enhanced public awareness of marine protection, prompting greater attention to environmental conservation measures. Impacts of Marine Environmental Issues: Respondents commonly believe that marine pollution has wide-ranging and profound impacts on ecosystems, society, and the economy. Marine pollution affects biodiversity and ecological balance and impacts economic activities such as fisheries and tourism, thereby affecting local and global economic sustainability. Moreover, resolving marine environmental issues is limited by the effectiveness of current environmental policies, requiring global collaborative efforts and sustained policy support. Respondents also mentioned that marine pollution could lead to food safety issues, such as the accumulation of heavy metals and harmful substances in seafood. These issues not only affect human health but could also cause social panic and affect social stability. Despite the public’s strong interest in marine pollution, specific actions toward marine protection are still insufficient. Most people’s marine conservation behaviors are limited to basic actions like avoiding littering in the ocean, and a deeper level of marine awareness and environmental behavior has not been fully developed. This reflects that China still needs to make further efforts to enhance public environmental awareness and behavior, especially in the field of marine protection.

*A11: “I think marine protection is very necessary. Like the nuclear wastewater, it has now flowed into our waters from Japan. Last year, when Japan discharged nuclear pollution, we asked our teachers whether it would affect humans because we work at sea and spend a lot of time there, wondering if it could impact us?”*

*A5: “If the ocean is polluted, our Earth’s water resources will also be polluted, and then it would affect the ecological cycle, which could have a significant impact on our planet. I think humanity should not overly explore the ocean, avoid damaging the ecosystem, and ensure that resource extraction does not destroy the ecology.”*

## Discussion

This study delves into the multidimensional structure, influencing factors, and role in promoting marine environmental behaviors of public ocean literacy. The results indicate that scientific knowledge, environmental ethics, conservation actions, policy understanding, and critical thinking together form the core elements of ocean literacy, aligning with the multidimensionality emphasized in existing research [7]. This study also reveals a widespread lack of marine knowledge among the Chinese public, consistent with previous findings in the Asian region, suggesting that the overall awareness of ocean literacy in China is low and requires systematic education and public engagement to enhance societal marine awareness [21, 28].

The findings demonstrate significant individual differences in ocean literacy levels, supporting the perspectives of the Value-Belief-Norm (VBN) theory that individual environmental beliefs and values profoundly affect their behaviors [12]. Specifically, individuals with high ocean literacy are more likely to exhibit proactive marine conservation behaviors, underscoring the important role of education and the cultivation of individual environmental values in enhancing ocean literacy levels. Therefore, this study emphasizes the importance of strengthening marine education to boost individual environmental responsibility and conservation behaviors.

The study also emphasizes the important roles of emotions and experiences in forming and enhancing ocean literacy. Interview results show that emotional connections to the ocean, such as awe, relaxation, and spiritual healing experiences, can promote individual attention to marine environmental conservation. This aligns with Brennan et, who argued that ocean literacy should encompass an emotional dimension [7]. Particularly, students in coastal areas, due to their daily direct contact with the ocean, are more likely to translate emotional attachment into actual conservation actions, a perspective also explained from the standpoint of psychoevolutionary theory [14]. Therefore, future marine education practices need to focus on integrating emotional experiences with knowledge education, creating more interactive and experiential learning opportunities to enhance public conservation awareness and behavioral intentions.

Marine environmental values and beliefs play a key role in shaping marine conservation behaviors among Chinese university students. The study reveals that most students hold strong environmental conservation values, with the formation of marine values significantly influenced by education, family environment, social networks, and geographical factors. Particularly, residents of coastal areas often have a higher sense of environmental responsibility, consistent with previous research findings that geographical location significantly affects individual environmental cognition and behavior [13, 43]. According to the Value-Belief-Norm (VBN) theory, when individuals possess strong marine conservation values, they develop a heightened environmental awareness and perceived responsibility, thereby more actively participating in marine conservation behaviors [17].

In the realm of marine policy and resource management, interviewees commonly agreed that challenges such as ocean pollution and resource overexploitation could be more effectively addressed through cross-regional and international collaboration, a perspective supported by existing international research [1]. However, interview data also indicated that the current implementation of marine resource management policies in China is insufficient, suggesting that future policy-making needs to focus more on the effectiveness of policy execution and broad public participation. Socio-cultural analysis showed that preserving marine cultural heritage has a positive impact on the formation of students’ sense of environmental responsibility. The research revealed that most students emphasized that protecting marine cultural heritage is not only a matter of cultural inheritance but also enhances individual environmental consciousness and social responsibility, aligning with the views of McKinley on the relationship between cultural heritage and environmental responsibility [44].

Additionally, this study pointed out that although sudden environmental incidents like nuclear wastewater discharges can temporarily heighten public environmental awareness, they are challenging to sustain in the long term. This result aligns with the perspectives in environmental behavior psychology, which state that sustained changes in environmental attitudes and behaviors require long-term, systematic education and policy interventions [11, 12]. Therefore, there is a need to develop and implement long-term effective public education programs and policy measures to continuously promote public conservation behavior. At the educational level, the interview results from this study highlighted the current limitations of marine education, namely that theoretical teaching is overly simplistic and lacks practical and interactive elements. This has resulted in students struggling to form a deep and lasting awareness of marine conservation. Respondents universally expressed a desire for enhanced marine awareness through interactive science communication platforms and practical experience activities, consistent with the perspectives on interactive marine education, which emphasize the importance of innovative educational forms in enhancing ocean literacy [41]. In summary, based on qualitative analysis and viewing from both international and local research perspectives, this study elucidates how marine literacy can promote university students’ environmental behaviors through mechanisms of environmental beliefs, values, and responsibility. These findings provide a theoretical and practical basis for further optimizing China’s marine education and enhancing public ocean literacy levels.

## Conclusion

This study employs qualitative methods to investigate how ocean literacy promotes marine conservation behaviors among Chinese university students. The findings reveal that the core components of ocean literacy—marine scientific knowledge, environmental ethics, conservation actions, policy understanding, and critical thinking—are crucial in cultivating and enhancing environmentally friendly behaviors. These components collectively shape a comprehensive understanding of the marine environment among individuals and ignite a willingness to actively participate in marine conservation.

The research highlights a general deficiency in marine knowledge among Chinese university students, particularly in terms of systematic depth, which hampers the development of their conservation behaviors. Consequently, strengthening marine education, especially in enhancing scientific literacy and critical thinking skills, is identified as key. Emotional connections and direct experiences also play an indispensable role in raising environmental awareness and fostering actual conservation behaviors. Additionally, effective marine policies and resource management, along with intensified international cooperation, are critical in addressing global marine environmental challenges.

The preservation and transmission of marine cultural heritage also demonstrate its importance in nurturing individuals’ sense of environmental responsibility. By enhancing education and public engagement through cultural and social dimensions, we can more effectively improve public ocean literacy and promote conservation behaviors. Finally, the study emphasizes that improvements in marine education should focus more on enhancing practical and interactive learning. Innovative educational methods, such as virtual reality technology and field-based experiential learning, can increase student engagement and deepen their understanding of marine issues. These findings provide specific guidelines for optimizing marine education strategies, enhancing public participation, and strengthening international cooperation, laying a theoretical and practical foundation for future marine conservation efforts.

## Acknowledgments

This project did not receive any funding.This work was supported by the School-level Education and Teaching Reform Project, Guangdong Ocean University, 2023 (Project Code: PX-972023240); 2024 Special Project on Mental Health in Higher Education Institutions sponsored by the Guangdong Provincial Association for the Study of Ideological and Political Education in Higher Education Institutions (Project No. 2024XLZX47); Counselor Zhang Li’s Studio at Guangdong Ocean University. We gratefully acknowledge the help of the above projects in this research.

## Author Contributions

Conceived and designed the experiments:Linzhao Wang, Bo Gao.

Methodology: Li Zhang, Xiangwei Chang.

Investigation: Bo Gao, Li Zhang.

Software: Linzhao Wang.

Visualization: Xiangwei Chang, Linzhao Wang.

Project administration: Linzhao Wang.

Writing – review & editing: Linzhao Wang, Bo Gao, Li Zhang.

## References

1. Masterson-Algar P, Jenkins SR, Windle G, Morris-Webb E, Takahashi CK, Rosa I, et al. When One Health Meets the United Nations Ocean Decade: Global Agendas as a Pathway to Promote Collaborative Interdisciplinary Research on Human-Nature Relationships. Front Psychol. 2022;13:809009.

2. Worm B, Barbier EB, Beaumont N, Duffy JE, Folke C, Halpern BS, et al. Impacts of biodiversity loss on ocean ecosystem services. Science. 2006;314(5800):787-90.

3. Claudet J, Bopp L, Cheung WWL, Devillers R, Escobar-Briones E, Haugan P, et al. A Roadmap for Using the UN Decade of Ocean Science for Sustainable Development in Support of Science, Policy, and Action. One Earth. 2020;2(1):34–42. doi: 10.1016/j.oneear.2019.10.012.

4. Ryabinin V, Barbière J, Haugan P, Kullenberg G, Smith N, McLean C, et al. The UN decade of ocean science for sustainable development. Front Mar Sci. 2019;6:470.

5. Cava F, Schoedinger S, Strang C, Tuddenham P. Science content and standards for ocean literacy: A report on ocean literacy. Potomac Falls, VA; 2005.

6. Fauville G, Strang C, Cannady MA, Chen Y-F. Development of the International Ocean Literacy Survey: measuring knowledge across the world. Environ Educ Res. 2018;25(2):238–63. doi: 10.1080/13504622.2018.1440381.

7. Brennan C, Ashley M, Molloy O. A System Dynamics Approach to Increasing Ocean Literacy. Front Mar Sci. 2019;6. doi: 10.3389/fmars.2019.00360.

8. McKinley E, Burdon D, Shellock RJ. The evolution of ocean literacy: A new framework for the United Nations Ocean Decade and beyond. Mar Pollut Bull. 2023;186:114467. Epub 2022/12/15. doi: 10.1016/j.marpolbul.2022.114467. PMID: 36516497.

9. Kelly R, Evans K, Alexander K, Bettiol S, Corney S, Cullen-Knox C, et al. Connecting to the oceans: supporting ocean literacy and public engagement. Rev Fish Biol Fish. 2022;32(1):123–43. Epub 2021/02/17. doi: 10.1007/s11160-020-09625-9. PMID: 33589856

10. Ashley M, Pahl S, Glegg G, Fletcher S. A Change of Mind: Applying Social and Behavioral Research Methods to the Assessment of the Effectiveness of Ocean Literacy Initiatives. Front Mar Sci. 2019;6. doi: 10.3389/fmars.2019.00288.

11. Ajzen I. The theory of planned behavior. Organ Behav Hum Decis Process. 1991;50(2):179–211.

12. Stern PC. New Environmental Theories: Toward a Coherent Theory of Environmentally Significant Behavior. J Soc Issues. 2000;56(3):407–24. doi: 10.1111/0022-4537.00175.

13. Schwartz SH. Normative influences on altruism. In: Berkowitz L, editor. Advances in Experimental Social Psychology. Vol 10. Elsevier; 1977. 221-79.

14. Longo SB, Clark B. An Ocean of Troubles: Advancing Marine Sociology. Soc Probl. 2016;63(4):463–79. doi: 10.1093/socpro/spw023.

15. Tsai L-T, Sasaki T, Wu C-K, Chang C-C. Ocean literacy among Taiwanese and Japanese high school students. Mar Policy. 2023;150. doi: 10.1016/j.marpol.2023.105555.

16. Fauville G. Digital technologies as support for learning about the marine environment: Steps toward ocean literacy. 2017.

17. Stern PC, Dietz T, Abel T, Guagnano GA, Kalof L. A value-belief-norm theory of support for social movements: The case of environmentalism. Hum Ecol Rev. 1999:81–97.

18. Schwartz SH, Bilsky W. Toward a universal psychological structure of human values. J Pers Soc Psychol. 1987;53(3):550.

19. Hilal AH, Alabri SS. Using NVivo for data analysis in qualitative research. Int Interdiscip J Educ. 2013;2(2):181–6.

20. McPherson K, Wright T, Tyedmers P. Examining the Nova Scotia Science Curriculum for International Ocean Literacy Principle Inclusion. Int J Learn Teach Educ Res. 2018;17(11):1–16. doi: 10.26803/ijlter.17.11.1.

21. Guest H, Lotze HK, Wallace D. Youth and the sea: Ocean literacy in Nova Scotia, Canada. Mar Policy. 2015;58:98–107. doi: 10.1016/j.marpol.2015.04.007.

22. Chang C-C. Development of Ocean Literacy Inventory for 16- to 18-Year-Old Students. SAGE Open. 2019;9(2). doi: 10.1177/2158244019844085.

23. Lee A. Analyzing the Effects of AI Education Program based on AI Tools. J-Institute. 2021.

24. Fauville G. Ocean Literacy in the Twenty-First Century. In: Exemplary Practices in Marine Science Education. 2019. 3-11.

25. Kollmuss A, Agyeman J. Mind the Gap: Why do people act environmentally and what are the barriers to pro-environmental behavior? Environ Educ Res. 2010;8(3):239–60. doi: 10.1080/13504620220145401.

26. De Leeuw A, Valois P, Ajzen I, Schmidt P. Using the theory of planned behavior to identify key beliefs underlying pro-environmental behavior in high-school students: Implications for educational interventions. J Environ Psychol. 2015;42:128–38. doi: 10.1016/j.jenvp.2015.03.005.

27. Charmaz K. Grounded theory. In Qualitative Psychology: A Practical Guide to Research Methods. 3rd ed. 2015. 53-84.

28. Wang Linzhao, Cho Jin-ho. Analysis of Differences in Marine Environmental Concerns and Ocean Literacy Levels Among Chinese University Students. Fisheries and Oceans Education and Research. 2024;36(3):510–24. doi: 10.13000/JFMSE.2024.6.36.3.510

29. Severin MI, Akpetou LK, Annasawmy P, Asuquo FE, Beckman F, Benomar M, et al. Impact of the citizen science project COLLECT on ocean literacy and well-being within a north/west African and south-east Asian context. Front Psychol. 2023;14:1130596. Epub 2023/06/30. doi: 10.3389/fpsyg.2023.1130596. PMID: 37388649

30. Liu GY, Lin YC, Yeh TK. Motivating Individuals to Take Responsible Ocean Action: The Mediatory Effects of Attitude toward the Ocean. Int J Environ Res Public Health. 2023;20(3). Epub 2023/02/12. doi: 10.3390/ijerph20032676. PubMed PMID: 36768042

31. Schwerdtner Manez K, Stoll-Kleemann S, Rozwadowski HM. Ocean literacies: the promise of regional approaches integrating ocean histories and psychologies. Front Mar Sci. 2023;10. doi: 10.3389/fmars.2023.1178061.

32. Hartig T, Mitchell R, de Vries S, Frumkin H. Nature and health. Annu Rev Public Health. 2014;35:207–28. Epub 2014/01/07. doi: 10.1146/annurev-publhealth-032013-182443. PMID: 24387090.

33. Von Schuckmann K, Holland E, Haugan P, Thomson P. Ocean science, data, and services for the UN 2030 Sustainable Development Goals. Mar Policy. 2020;121:104154.

34. Hu YX, Li ST, Chen RR, Tian H. Progress of research on pro-environmental behavior in the past 20 years. Psychological Research. 2021;14(5):428.

35. Mokos M, De-Bastos E, Realdon G, Wojcieszek D, Papathanasiou M, Tuddenham P. Navigating Ocean Literacy in Europe: 10 years of history and future perspectives. Mediterr Mar Sci. 2022. doi: 10.12681/mms.26989.

36. Zielinski T, Kotynska-Zielinska I, Garcia-Soto C. A Blueprint for Ocean Literacy: EU4Ocean. Sustainability. 2022;14(2). doi: 10.3390/su14020926.

37. Brymer E, Freeman E, Richardson M. Editorial: One Health: The Well-being Impacts of Human-Nature Relationships. Front Psychol. 2019;10:1611. Epub 2019/09/05. doi: 10.3389/fpsyg.2019.01611. PubMed PMID: 31481906

38. McKinley E, Acott T, Yates KL. Marine social sciences: Looking towards a sustainable future. Environ Sci Policy. 2020;108:85–92. doi: 10.1016/j.envsci.2020.03.015.

39. Qin H, Gao YH. On the environmentally friendly behaviors of island residents and their influencing factors. china population, resources and environment. 2020;30(4):125-35.

40. Fauville G, Dupont S, von Thun S, Lundin J. Can Facebook be used to increase scientific literacy? A case study of the Monterey Bay Aquarium Research Institute Facebook page and ocean literacy. Comput Educ. 2015;82:60–73.

41. Dupont S, Fauville G. Ocean literacy as a key toward sustainable development and ocean governance. In: Handbook on the Economics and Management of Sustainable Oceans. Edward Elgar Publishing; 2017. 519-37.

42. Kelly R, Elsler LG, Polejack A, van der Linden S, Tönnesson K, Schoedinger SE, et al. Empowering young people with climate and ocean science: Five strategies for adults to consider. One Earth. 2022;5(8):861–74. doi: 10.1016/j.oneear.2022.07.007.

43. Worm B, Elliff C, Fonseca JG, Gell FR, Serra-Gonçalves C, Helder NK, et al. Making ocean literacy inclusive and accessible. Ethics Sci Environ Polit. 2021;21:1–9. doi: 10.3354/esep00196.

44. McKinley E, Burdon D. Understanding Ocean Literacy and Ocean Climate-Related Behaviour Change in the UK-Work Package 1: Evidence Synthesis. Hull: Daryl Burdon Ltd; 2020. Available online at: https://darylburdon.co.uk.

